# 4E-BP2-dependent translational control in GABAergic interneurons is required for long-term memory

**DOI:** 10.1101/2025.10.14.682450

**Authors:** Ziying Huang, Niaz Mahmood, Konstantina Psycharis, Kevin Lister, Mehdi Hooshmandi, Nikhil Inturi, Diana Tavares-Ferreira, Shane Wiebe, Arkady Khoutorsky, Nahum Sonenberg

**Author notes:** **Corresponding author**: Nahum Sonenberg, McGill University, Department of Biochemistry, Goodman Cancer Research Institute, 1160 Pine Avenue West, Montréal, QC, Canada, H3A 1A3.

## Abstract

mRNA translational repression by eukaryotic initiation factor 4E-binding proteins (4E-BPs), plays a critical role in synaptic plasticity and the formation of long-term memory (LTM). Among the three 4E-BP paralogs, 4E-BP2 is the predominant form expressed in neurons, and its full-body deletion in mice causes memory deficits. Mice lacking 4E-BP2 in GABAergic inhibitory interneurons, but not excitatory neurons, display autistic-like behaviors and deficits in object location and recognition. The specific mRNAs translationally regulated by 4E-BP2 in GABAergic interneurons, and how they contribute to spatial and associative memory, are unknown. Here, we show that conditional knockout (cKO) mice lacking 4E-BP2 selectively in GABAergic interneurons exhibit impairments in long-term spatial and contextual fear memory formation. We further demonstrate that 4E-BP2 deletion controls the translation of selective mRNAs in interneurons without increasing general protein synthesis. One of the mRNAs is *Gal*, which encodes a neuropeptide that modulates memory. Our findings provide evidence that 4E-BP2 selectively controls the translation of a subset of mRNAs in inhibitory neurons that are required for LTM formation.

## Introduction

Memory is a fundamental cognitive process that encompasses the encoding, storage, and retrieval of information. The formation of long-term memory (LTM) relies on synaptic plasticity and neuronal *de novo* protein synthesis (Casadio et al., 1999; Goelet et al., 1986; Kandel, 2001; Klann & Sweatt, 2008). Eukaryotic mRNAs contain a cap structure (cap, m7GpppN, where N is any nucleotide) at the 5’ terminus (Shatkin, 1976). mRNA translation commences when the mRNA 5’ cap-binding protein complex, eIF4F, binds to the cap. eIF4F is composed of 3 subunits, eIF4E, the cap binding protein; eIF4A, an RNA helicase; and eIF4G, a scaffolding protein (Amiri et al., 2025; Pelletier et al., 2015). The eIF4E-binding proteins (4E-BPs) repress cap-dependent translation initiation by binding to eIF4E and thus interdicting eIF4E-eIF4G interaction and eIF4F complex formation (Gingras et al., 1999; Lin et al., 1994; Mader et al., 1995; Marcotrigiano et al., 1999; Pause et al., 1994; Poulin et al., 1998; Sonenberg & Dever, 2003). Hierarchical phosphorylation of 4E-BPs by the mechanistic/mammalian target of rapamycin complex 1 (mTORC1) facilitates their dissociation from eIF4E, allowing the assembly of the eIF4F complex (Bah et al., 2015; Gingras et al., 2001; Lin et al., 1994; Matsuo et al., 1997; Pause et al., 1994; Sonenberg & Gingras, 1998). Among the three 4E-BP paralogs, 4E-BP2 is the predominant paralog in neurons (A. Aguilar-Valles et al., 2021; Banko et al., 2005). 4E-BP2 is crucial for translation of synaptic mRNAs, many of them containing 5′ terminal oligopyrimidines (5’TOP) (Gkogkas et al., 2013; Hochstoeger & Chao, 2024; Hochstoeger et al., 2024; Nygård & Nilsson, 1990; Ran et al., 2013; Tang et al., 2001).

Full-body deletion of 4E-BP2 mimics mTORC1 hyperactivation by constitutively relieving eIF4E inhibition. Deletion of 4E-BP2 lowers the threshold for early-phase long-term potentiation (E-LTP) in hippocampus but disrupt the maintenance of late-phase long-term potentiation (L-LTP) and leads to exaggerated mGluR-dependent long-term depression (mGluR-LTD) (Banko et al., 2005; Ran et al., 2009; Stoica et al., 2011). These synaptic alterations are accompanied by impairments in LTM formation, including associative, spatial, recognition, and working memory, which are thought to be due to abnormal synaptic plasticity (Banko et al., 2007; Banko et al., 2005). While most research has focused on excitatory neurons, more recent studies highlight a critical role of mTORC1 signaling and eIF4E-dependent translational control in inhibitory interneurons in modulating neuronal circuits and information processing during learning (Artinian et al., 2019; Shrestha, Ayata, et al., 2020; Shrestha, Shan, et al., 2020; Tajima et al., 2023). We recently demonstrated that GABAergic inhibitory, but not excitatory, neuron-specific deletion of 4E-BP2 recapitulates most of the behavioral phenotypes observed in global 4E-BP2 knockout (KO) mice, including autistic-like behaviors and recognition memory impairments (Huang et al., 2024; Wiebe et al., 2019). These findings indicate that mRNA translational control by 4E-BP2 in GABAergic interneurons is indispensable for LTM formation and neurodevelopment (Huang et al., 2024; Wiebe et al., 2019). However, the specific mRNAs translationally regulated by 4E-BP2 in GABAergic neurons and the impact of this regulation on long-term fear associative memory and spatial memory are not known.

Here, we employed a combination of genetic, molecular, and behavioral approaches to elucidate the role of 4E-BP2 in inhibitory neurons during memory formation. Using a conditional knockout mouse model (*Eif4ebp2^flx/flx^:Gad2-Cre*), we assessed memory formation following contextual fear conditioning (CFC) and the Morris water maze (MWM) tests, two widely used hippocampus-dependent memory tasks (Fanselow, 2000; Logue et al., 1997; Remaud et al., 2014; Stackman et al., 2016). Mechanistically, the mTORC1 signaling pathway plays a critical role in promoting LTM formation in the latter behavior paradigms (Artinian et al., 2019; MacCallum & Blundell, 2020). Our results reveal that 4E-BP2 deletion in GABAergic neurons is sufficient to impair hippocampus-dependent LTM formation. We assessed global protein synthesis in inhibitory neurons using fluorescent non-canonical amino-acid tagging (FUNCAT) and uncovered ribosome-associated mRNAs using viral translating ribosome affinity purification (vTRAP) sequencing. In GABAergic interneurons, 4E-BP2 deletion selectively affects the translation of specific mRNAs without altering general protein synthesis. One of the validated downregulated mRNAs encodes the neuropeptide galanin—a critical modulator of cognitive function (Beltran-Casanueva et al., 2024; Elliott-Hunt et al., 2004; Miller, 1998; Ögren et al., 2010). These findings underscore the importance of interneuron-specific translational control in memory formation and provide novel insights into how 4E-BP2 impacts the translatome of inhibitory circuits.

## Results

### 4E-BP2 deletion in GABAergic neurons impairs hippocampus-dependent long-term memory

Disruption of the mTORC1–4E-BP2 axis in GABAergic neurons perturbs cognitive function and neurodevelopment (Huang et al., 2025; Huang et al., 2024). To investigate the impact of 4E-BP2 deletion on LTM and translation of its target mRNAs, we utilized a conditional knockout (cKO) mouse model (*Eif4ebp2^flx/flx^:Gad2-Cre*) by selectively ablating 4E-BP2 in GABAergic neurons (Wiebe et al., 2019). Immunofluorescence analysis confirmed deletion of 4E-BP2 in GABAergic cells by co-staining for antibodies against 4E-BP2 and glutamate decarboxylase 67 (Gad67), a marker of GABAergic neurons, in cKO hippocampus (Fig. 1A and B) [86.86 ± 5.737% decrease relative to wild-type controls (control: 100 ± 5.467, n = 11 cells/mouse, 5 mice; cKO: 13.01 ± 1.740, n = 9–11 cells/mouse, 5 mice); *t*(4.802) = 15.14, *p* < 0.01; unpaired *t*-test with Welch’s correction].

**Figure 1.**
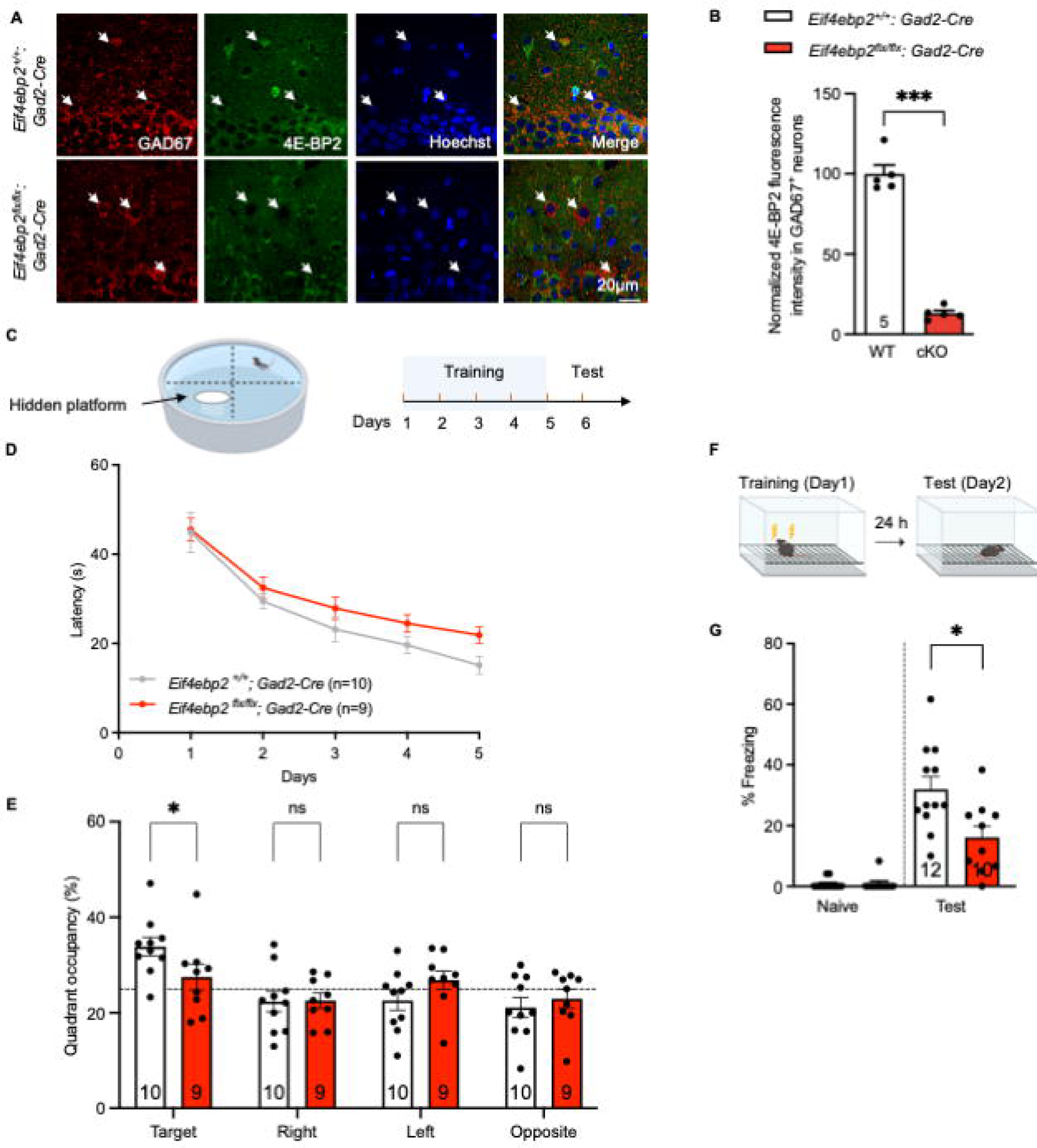
Conditional deletion of 4E-BP2 in GAD67+ inhibitory neurons impairs long-term memory. (A) Representative confocal images showing immunofluorescence staining of 4E-BP2 (green) in GAD67+ cells (red) from hippocampal sections of *Eif4ebp2^+/+^:Gad2-Cre* (WT) versus *Eif4ebp2^flx/flx^:Gad2-Cre* (cKO) mice. Hoechst (blue) labels nuclei. Scale bar, 20 μm; magnification 60×. (B) Quantification of mean integrated density of 4E-BP2 signal in GAD67+ neurons (9–11 cells per mouse; n = 5 mice per group). (C) Schematic diagram of the MWM training and testing protocol. (D) Spatial learning and memory were assessed by latency to locate the hidden platform across five consecutive acquisition days. (E) Probe test results, expressed as the percentage of time spent in the target quadrant on the final test day, demonstrated spatial memory impairment in 4E-BP2 cKO mice compared to WT. (F) Diagram of the weak contextual fear conditioning (CFC) paradigm (2-foot shocks, 0.7 mA, 2 s duration). (G) LTM assessed as freezing behavior during a 5-minute test, 24 hrs post-training. Behavioral experiments were conducted in the same cohort of mice: control (n = 10) vs. 4E-BP2 cKO (n = 9). Data represent individual mice and are expressed as mean ± SEM. *P < 0.05, ***P < 0.001, n.s., not significant.

To assess spatial memory, we subjected 4E-BP2 cKO mice to the MWM test, a hippocampus-dependent memory task (Fig. 1C) (Mahmood et al., 2024; Morris, 1981). During the five days of memory acquisition, both WT and cKO showed decreased escape latencies, indicating effective learning of the platform’s spatial location. However, cKO mice show overall higher escape latencies compared to WT (Fig. 1D). On the 5^th^ training day, cKO mice exhibited a trend of prolonged escape latency relative to WT, although this difference did not reach statistical significance [Fig. 1D; escape latency on day five (control: 15.10 ± 1.988 s, n = 10; cKO: 21.89 ± 1.831 s, n = 9; p = 0.065; two-way ANOVA)]. In the probe trial conducted 24 hours after the last training session (with the platform removed), cKO mice spent less time in the target quadrant than controls, indicating deficits in spatial memory (Fig. 1E; control: 33.86 ± 1.960 s, n = 10; cKO: 27.50 ± 2.686 s, n = 9; p = 0.0344; two-way ANOVA with Tukey’s post hoc test).

Hippocampus-dependent long-term associative learning was further studied using the CFC paradigm (Fanselow, 2000; Mahmood et al., 2024) (Fig. 1F). Control and cKO groups received foot-shock treatment in identical context boxes, and LTM was tested 24 hours later. 4E-BP2-Gad2 cKO mice demonstrated 1.97-fold reduced freezing behavior relative to controls during the LTM test, indicating impaired contextual fear memory (Fig. 1G; control: 31.94 % ± 4.107, n = 12; cKO: 16.17% ± 3.719, n = 10; *t*(20) = 2.797, *p* = 0.0111; unpaired *t*-test). Notably, deletion of 4E-BP2 in excitatory neurons does not affect contextual fear-associated memory (Fig. S1; control: 32.86% ± 4.724, n = 7; cKO: 37.62% ± 1.402, n = 7; *t*(7.05) = 0.9664, *p* = 0.3658; unpaired *t*-test with Welch’s t-test).

Taken together, the results demonstrate that 4E-BP2 function in GABAergic neurons is essential for hippocampus-dependent LTM, supporting a memory-promoting role for 4E-BP2-mediated translational regulation within inhibitory neurons.

### GABAergic neuron-specific 4E-BP2 deletion does not impact global protein synthesis and synaptic protein expression

To determine whether 4E-BP2-Gad2 cKO disrupts global protein synthesis in inhibitory neurons, we employed FUNCAT for labeling newly synthesized proteins *in situ* (Hooshmandi et al., 2024; Tom Dieck et al., 2012). Mice were fed a methionine-deficient diet and subsequently administered azidohomoalanine (AHA), a methionine analog that is incorporated into newly synthesized proteins (Fig. 2A). AHA-labeled proteins were covalently conjugated *via* click chemistry to a fluorescent dye that incorporates an alkyne functional group. The alkyne group on the dye reacts with the azide on AHA, allowing the newly synthesized proteins to be visualized by confocal microscopy (Fig. 2B). Quantitative analysis of AHA signal in GABAergic hippocampal neurons, identified by GAD67 immunoreactivity, indicated no difference in AHA incorporation between 4E-BP2 cKO mice and wild-type controls (Fig. 2C; control: 100 ± 2.520; 4E-BP2 cKO: 108.9 ± 9.020; *t*(4.620) = 0.9504, *p* = 0.3889; unpaired *t*-test with Welch’s correction). The results demonstrate that global protein synthesis in GABAergic neurons remained unchanged following 4E-BP2 deletion. This finding is consistent with observations from other *in vivo* and *in vitro* models, including full-body 4E-BP2 KO mice, 4E-BP1/2 double KO mouse embryonic fibroblasts, and Purkinje cell-specific 4E-BP2 cKO mice, which reported unaltered global protein synthesis in the absence of 4E-BP2 (Alasad et al., 2020; Banko et al., 2005; Dowling et al., 2010; Hooshmandi et al., 2021).

**Figure 2.**
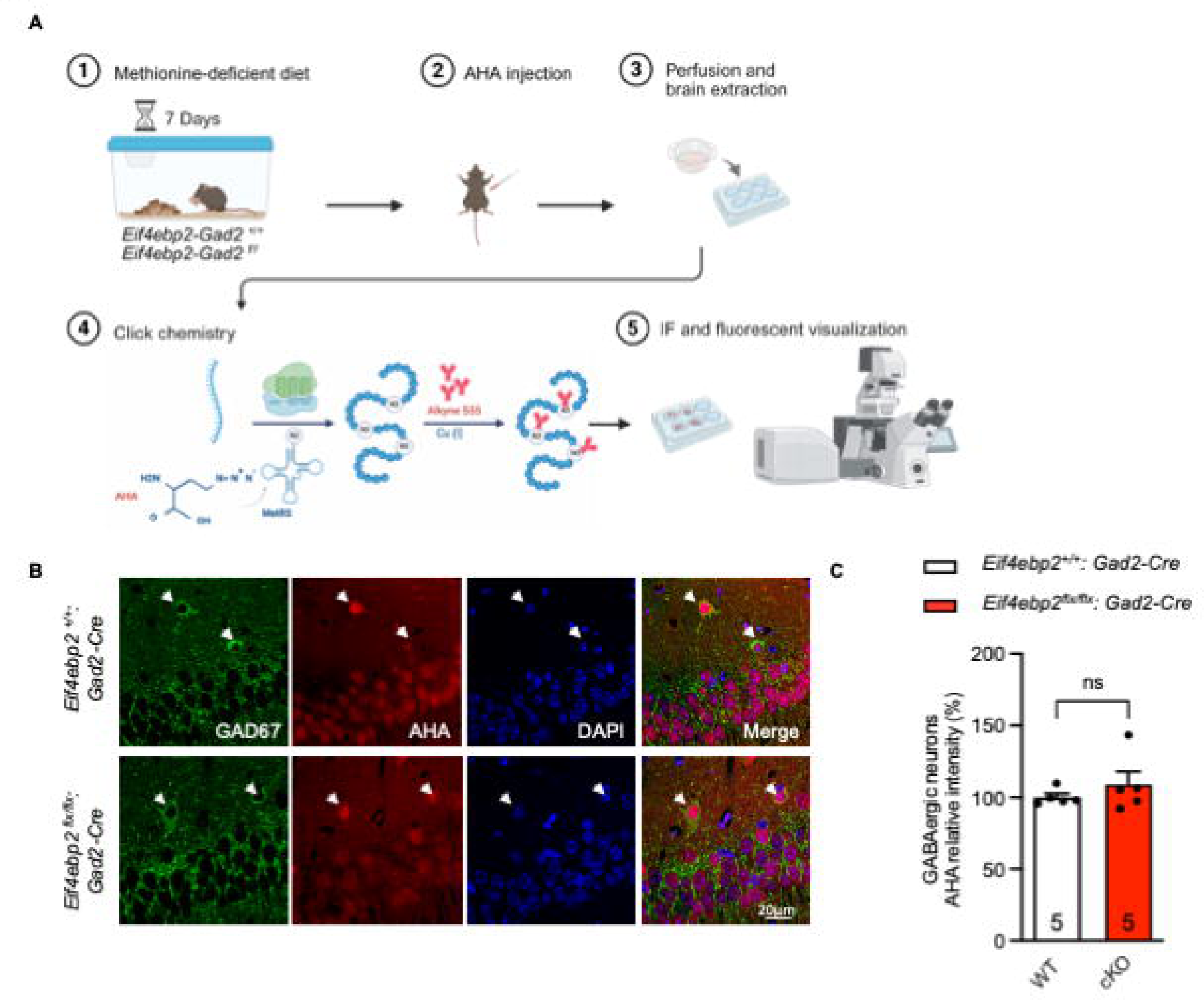
4E-BP2 deletion does not affect bulk protein synthesis in GABAergic neurons. (A) Schematic overview of the FUNCAT protocol used to label newly synthesized proteins. (B) Representative confocal images of AHA incorporation (red) in hippocampal GAD67+ neurons (green) in control *Eif4ebp2^+/+^:Gad2-Cre* (WT) versus *Eif4ebp2^flx/flx^:Gad2-Cre* (cKO) mice. Hoechst (blue) labels nuclei. Scale bar, 20 μm; magnification 60×. (C) Quantification of mean integrated density of AHA signal per cell (9–11 cells per mouse; n = 5 mice per group). Data represent individual mice and are presented as mean ± SEM. Statistical significance was determined using unpaired Student’s t-test. n.s., not significant.

To determine whether 4E-BP2 selectively controls the translation of mRNAs encoding inhibitory neuron-specific synaptic proteins, we performed Western Blot analysis on crude hippocampal lysates and olfactory bulb lysates (Fig. S2A, S2C). The olfactory bulbs were examined because of its high percentage (∼70%-80%) of GABAergic neurons (Bagley et al., 2007; Burton, 2017; Parrish-Aungst et al., 2007). Protein levels of the inhibitory markers vesicular GABA transporter (VGAT), GABA B receptor 2 (GABABR2), gephyrin, as well as the AMPAR subunit glutamate receptor 1 (GluR1) were assessed by Western blot. Our results show no changes in the expression levels of these synaptic markers between the control and 4E-BP2 cKO mice (Fig. S2A-D). Levels of neuroligin 2 (NLGN2), which is enriched in inhibitory synapses and plays a critical role in the excitation-inhibition balance, were not changed, either (Fig. S2A-D) (Ali et al., 2020; Fu & Vicini, 2009; Gkogkas et al., 2013). Thus, the data demonstrate that deletion of 4E-BP2 does not alter protein amounts of key synaptic proteins.

### Translational landscape alteration in GABAergic interneurons lacking 4E-BP2

To identified which mRNAs are impacted by 4E-BP2 deletion, we utilized vTRAP, a technique that enables cell type-specific isolation of actively translating mRNAs (Nectow et al., 2017). We injected adeno-associated virus AAV5-DIO-eGFP-L10a bilaterally into the dorsal hippocampus of control (Gad2-Cre) and 4E-BP2 cKO mice to drive Cre-dependent expression of enhanced green fluorescent protein (eGFP)-tagged ribosomal protein L10a, selectively in GABAergic neurons (Fig. 3A). Co-localization of GAD67 and eGFP signal confirmed the specific expression of eGFP-L10a in inhibitory neurons (Fig. 3B). eGFP-immunoprecipitated (IP) ribosomes were isolated from dorsal hippocampal tissue (pooled from 3 mice/replicate, n = 4 replicates/condition), and the associated mRNAs were sequenced along the bulk input (IN) RNA fractions.

**Figure 3.**
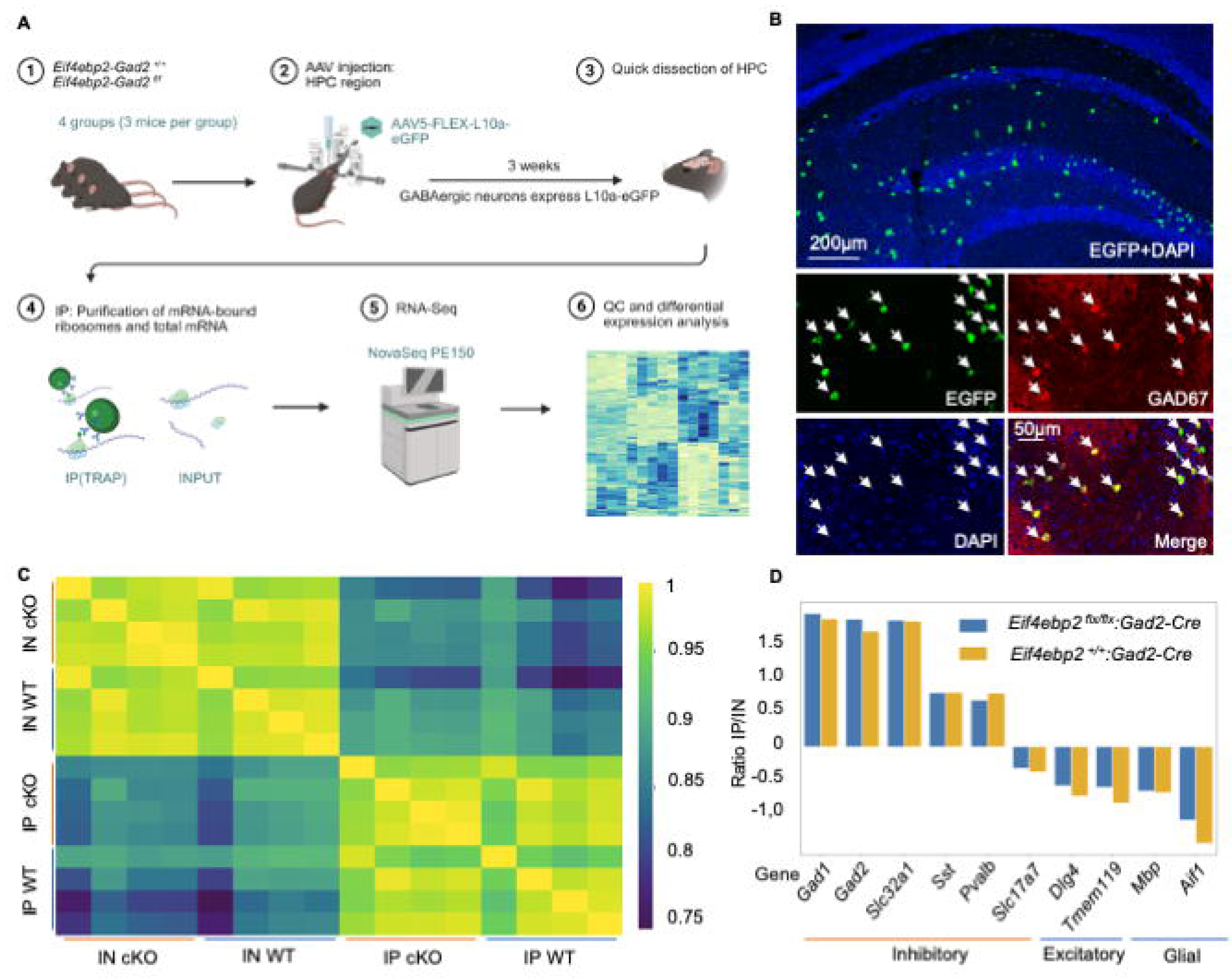
vTRAP-Seq enables cell-type-specific profiling of translational changes in hippocampal inhibitory neurons. (A) Schematic of the vTRAP strategy used to isolate ribosome-bound mRNAs from GABAergic neurons. (B) Immunofluorescence images showing selective expression of eGFP-L10a (green) in GAD67+ expressing neurons (red) following AAV-eGFP-L10a injection into Gad2-Cre mice. Hoechst (blue) stains nuclei. Scale bars: 500 μm (upper panel), 50 μm (lower panels). (C) Heatmap of correlation coefficients showing distinct clustering between input (IN) and IP samples. Samples are clearly separated by condition (control vs. cKO), demonstrating specificity of vTRAP-seq. (D) Validation of vTRAP enrichment: quantitative sequencing data from immunoprecipitated (IP) fractions confirms selective enrichment for GABAergic markers (*e.g*., *Gad1*, *Gad2, Slc32a1, Sst, Pvalb*) and depletion of excitatory and glial markers (*e.g*., *Slc17a7*, Dig4, Tmem119, Mbp, *Aif1*).

Quality control metrics showed high reproducibility across biological replicates: Pearson correlation coefficients for transcript-per-million (TPM) expression ranged from 0.98 to 1.0 (Fig. S3A). Hierarchical clustering and heatmap visualization of TPM values revealed clear separation between IP and IN fractions, confirming successful enrichment of ribosome-bound mRNAs (Fig. 3C; S3B). Quantile normalization was applied to correct for sequencing depth and ensure uniform expression scaling across samples (Fig. S3C). Marker gene analysis validated the cell type-specificity: the immunoprecipitated (IP) fraction was enriched for mRNAs encoding canonical inhibitory neuron markers, including *Gad1* (glutamate decarboxylase 1), *Gad2* (glutamate decarboxylase 2), *Slc32a1* (vesicular inhibitory amino acid transporter), *Sst* (somatostatin), and *Pvalb* (parvalbumin). Whereas mRNAs specific to excitatory neurons, such as *Slc17a7* (vesicular glutamate transporter 1) and *Dlg4* (discs large MAGUK scaffold protein 4), and glial cells, including *Tmem119* (transmembrane protein 119), *Mbp* (myelin basic protein), and *Aif1* (allograft inflammatory factor 1/Iba1), were depleted.

Comparative analysis of ribosome-bound transcripts from control versus 4E-BP2 cKO mice revealed 92 differentially translated mRNAs (Fig. 4A). Of these, 41 were upregulated and 51 were downregulated in the cKO mice compared to WT controls. A heatmap of correlation coefficients across genotypes indicated genotype-specific divergence in translational profiles (Fig. 4A, S4A). 4E-BP2 depletion is generally expected to augment translation of mRNAs because 4E-BPs are suppressors of translation initiation. However, transcriptome-wide studies reported both stimulation and inhibition of translation in cells (Amorim et al., 2018; Cridge et al., 2010). These findings suggest that effects of 4E-BP2 depletion on translation may be context-dependent and the observed activation of mRNA translation is likely indirect and not understood. Among the top 15 differentially translated mRNAs (Fig. 4A, S4B), *Gal*, *Th* (Tyrosine Hydroxylase), *ccl21* (C-C motif chemokine ligand 21), and *Ccr5* (C-C chemokine receptor type 5 are associated with aging-related memory deficits and synaptic regulation,) (Maynard et al., 2020; Robinson & Crawley, 1993; Zhou et al., 2016). We could not validate TH, CCL21 and CCR5. *Gal* (galanin) was among the most significantly downregulated (3.42-fold) mRNAs in the 4E-BP2 cKO IP fraction (Fig. 4A). Galanin is expressed in hippocampal and hypothalamic GABAergic neurons and plays an important role in modulating learning and memory through its inhibitory effects on excitatory neurotransmission and by modulating neuronal excitability (Elliott-Hunt et al., 2004; Keimpema et al., 2014; Kroeger et al., 2018; Massey et al., 2003; Qualls-Creekmore et al., 2017; Wrenn et al., 2004). Given its established role as a Gi-coupled neuropeptide receptor ligand that acts as a neuroprotective factor in the hippocampus, reduced *Gal* mRNA translation may contribute to the observed memory deficits (Beltran-Casanueva et al., 2024; Elliott-Hunt et al., 2004; Yoshitake et al., 2011). Previous publications and Ribo-Seq datasets of neuropil versus somatic compartments indicate that galanin is preferentially translated locally in neurites, predominantly in non-excitatory neurons (Fig. S4C) (Glock et al., 2021; Vila-Porcile et al., 2009). Immunofluorescent co-labeling of galanin and GAD67 revealed a 28.57% decrease in galanin immunoreactivity in the neurites of GABAergic neurons relative to controls (Fig. 4C; 4D; control: 1 ± 0.096, n = 1-4 neurites/mouse, 6 mice; cKO: 0.714 ± 0.032, n = 1-4 neurites/mouse, 6 mice; t(20)=3.016, p = 0.0136; two-way ANOVA). However, galanin expression within the cell body of the inhibitory neuron remained largely unchanged in cKO mice compared to controls (Fig. 4B; 4D; control: 1 ± 0.063, n = 5-7 cells/mouse, 6 mice; cKO: 0.884 ± 0.062, n = 5-7 cells/mouse, 6 mice; t(20)=1.223, p = 0.4158; two-way ANOVA), indicating a compartment-specific dysregulation of galanin resulting from the deletion of 4E-BP2.

**Figure 4.**
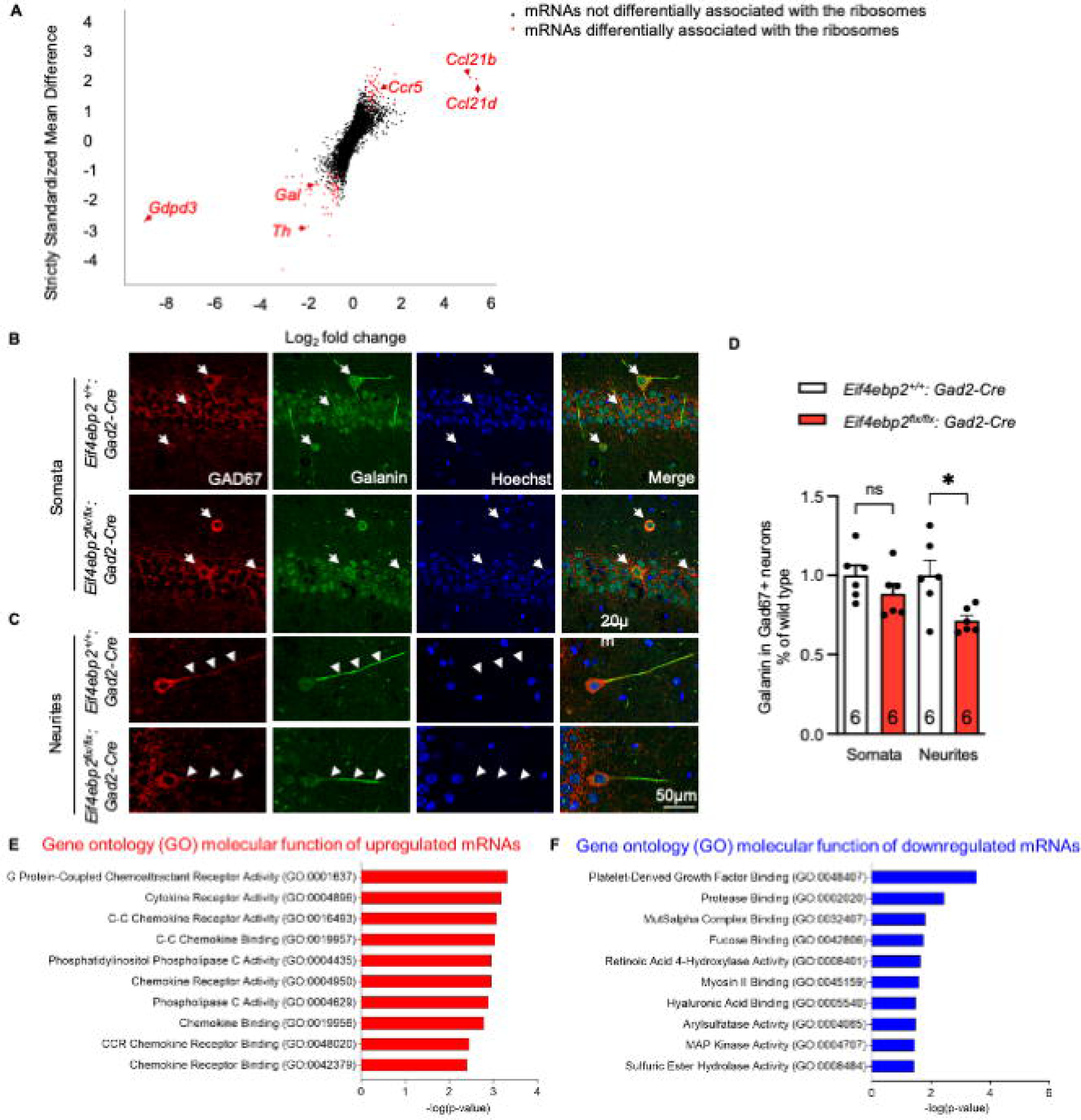
vTRAP-Seq reveals that loss of 4E-BP2 alters translation of specific mRNAs in hippocampal inhibitory neurons. (**A)** Dual flashlight plot showing log_2_ fold change (Log_2_FC) versus strictly standardized mean difference (SSMD) for translated mRNAs in IP samples. Red dots indicate significantly differentially translated mRNAs in control *Eif4ebp2^+/+^:Gad2-Cre* (WT) versus *Eif4ebp2^flx/flx^:Gad2-Cre* (cKO) hippocampal GABAergic neurons: upregulated (SSMD ≥ 0.9, BC ≥ 0.5, FC ≥ 1.33) and downregulated (SSMD ≤ -0.9, BC ≤ -0.5, FC ≤ 0.75). (B-C) Representative confocal images of Galanin protein expression (Green) in hippocampal GAD67+ neurons (red) in control and 4E-BP2 cKO mice. Scale bar, 20 μm; magnification 60×. Hoechst (blue) stains nuclei. (D) Quantification of galanin expression in GABAergic neurons’ somata and neurites (1–7 neurons per mouse, n = 6 mice per group). (E-F) GO pathway enrichment analysis of 92 differentially translated mRNAs in 4E-BP2-deficient GABAergic neurons. Statistical significance was determined using two-way ANOVA. *P < 0.05, n.s., not significant. Data represent individual mice and are expressed as mean ± SEM.

Gene ontology (GO) analysis revealed that mRNAs upregulated in 4E-BP2 cKOs were linked to chemokine receptor activity, indicating disruption of neuromodulators and intracellular signaling critical for memory (Fig. 4E)(Oppermann, 2004; Zhou et al., 2016). In contrast, downregulated transcripts were enriched for those encoding Platelet-Derived growth factor binding and protease binding proteins, which are critical for spine morphology and memory formation (Fig. 4F)(Funa & Sasahara, 2014; Shioda et al., 2012). The findings reveal that 4E-BP2 governs a specialized translational program in GABAergic neurons, regulating a subset of mRNAs which function in synaptic structure and neuromodulatory signaling, including the neuropeptide mRNA *Gal*. Regulating a subset of mRNAs involved in synaptic structure and neuromodulatory signaling, including the neuropeptide gene Gal. Rather than broadly suppressing protein synthesis, 4E-BP2 controls the translation of specific mRNAs, which impacts memory and behavior.

## Discussion

In this study, we investigated the cell type-specific role of the translational repressor 4E-BP2 within inhibitory neurons and its contribution to hippocampus-dependent LTM. Selective deletion of 4E-BP2 in GABAergic interneurons impaired spatial and contextual fear memory. 4E-BP2 deletion in inhibitory neurons changed ribosome-association with a subset of mRNAs without inducing global alterations in protein synthesis.

Rapid, activity-induced protein synthesis, particularly through eIF4E-mediated cap-dependent translation, is indispensable for memory consolidation (Costa-Mattioli et al., 2009; Gkogkas et al., 2010; Stoica et al., 2011). Full-body 4E-BP2 KO mice exhibit deficits in LTM tasks (Banko et al., 2007; Banko et al., 2005; Ran et al., 2013). We show that GABAergic-specific 4E-BP2 deletion is sufficient to cause hippocampus-dependent LTM deficits, highlighting a critical role for inhibitory neuron translation. We previously reported that selective disruption of the mTORC1–4E-BP2 axis, either by deleting *RPTOR* or *4E-BP2,* or through chemogenetic manipulations, in GABAergic neurons, but not in excitatory neurons, impairs object location and object recognition tasks (Huang et al., 2025; Huang et al., 2024). Our current findings address a knowledge gap by linking 4E-BP2 and translation in GABAergic interneurons to two key LTM paradigms—MWM and CFC. The importance of 4E-BP2’s upstream mTORC1 pathway in somatostatin expressing (SST^+^) interneurons for associative memory formation has also been shown (Artinian et al., 2019; Honoré et al., 2022; Honoré & Lacaille, 2022). Also, 4E-BP2 cKO in Purkinje cells, the cerebellum’s principal inhibitory neurons, impairs spatial memory performance in MWM (Hooshmandi et al., 2021). Taken together, these findings reinforce the role of 4E-BP2–mediated translational control across inhibitory neuron subtypes in regulating memory.

While 4E-BP2 deletion in excitatory neurons does not affect contextual fear memory, it is required in amygdala excitatory neurons for cue fear conditioning (Alapin et al., 2023). These findings indicate region and cell type-specific roles for translational control in memory: 4E-BP2 is essential in hippocampal inhibitory neurons for contextual memory, but in the amygdala excitatory neurons for cue-based learning. While 4E-BP2 in excitatory neurons appears dispensable for a few forms of hippocampus-dependent LTM, translational regulation in these neurons remains crucial for certain hippocampus-dependent memory. This is consistent with extensive proteome remodeling in both hippocampal excitatory neurons and SST^+^ interneurons following CFC (Oliveira et al., 2025). The eIF2α-dependent integrated stress response in excitatory neurons also contribute to memory formation by selectively regulating translation of specific mRNA subsets (e.g., *Atf4,* Activating Transcription Factor 4) (Mahmood et al., 2024; Sharma et al., 2020; Simbriger et al., 2021). Overall, a large body of evidence supports circuit-specific and cell type-specific translational control mechanisms for distinct memory phenotypes.

We conclude that 4E-BP2 does not globally enhance protein synthesis but rather selectively controls translation of specific mRNAs that are particularly sensitive to eIF4E activity (Alasad et al., 2020; Banko et al., 2005; Dowling et al., 2010; Hooshmandi et al., 2021). vTRAP profiling identified 92 differentially translated mRNAs in 4E-BP2 KO GABAergic neurons—a number similar to that seen in eIF4E overexpression models in non-neuronal cells (Ghram et al., 2023). Unlike prior studies in full-body 4E-BP2 KO model, we did not detect neuroligin mRNAs within the GABAergic neuron IP fractions, possibly reflecting cell-type differences (Gkogkas et al., 2013; Glock et al., 2021; Suresh et al., 2025). Similar to our findings, Cridge et al. (2010) demonstrated that 4El1BPs in yeast modulate translation of over 1,000 mRNAs, with many being both upl1 and downl1regulated in mutants lacking specific 4El1BPs (Caf20p and Eap1p) (Cridge et al., 2010). Amorim et al. (2018) also reported that eIF4El1dependent translation is bidirectional, showing selective increases and decreases in translation among different mRNAs in the nervous system (Amorim et al., 2018). In our case, deleting the translational repressor 4E-BP2 increased L10a-associated binding for a subset of mRNAs, including *Ccl21* and *Ccr5*. CCL21 is a homeostatic chemokine that is expressed in damaged neurons, rather than healthy neurons, where it’s up-regulation promotes neuroinflammation (Biber & Boddeke, 2014; Biber et al., 2011; Fülle et al., 2018; van Weering et al., 2010). CCR5, a β-chemokine receptor, has been shown to supress cortical plasticity and hippocampus-dependent memory when overexpressed in neurons (Zhou et al., 2016). Increased translation engagement of mRNAs related to chemokine receptor activity and chemokine binding activity may contribute to the memory and autism-like phenotypes in our 4E-BP2 cKO model. We also observed reduced ribosomal association of certain mRNAs, notably *Gal* mRNA. This effect may reflect competitions among mRNAs for translation initiation factors or unknown feedback regulations. Given the known preference of eIF4E for mRNAs with structured 5’ untranslated regions (UTRs) or terminal oligopyrimidine (TOP) motif, ribosomes may become occupied with such mRNAs, reducing access of others (such as *Gal*) to the translation machinery (Hochstoeger & Chao, 2024; Koromilas et al., 1992; Thoreen et al., 2012; Truitt et al., 2015). However, the exact mechanism behind this is still unknown and should be explored in future research.Galanin, an inhibitory neuropeptide is translated locally in neurites in several brain regions, including hypothalamus and forebrain (Glock et al., 2021; Keimpema et al., 2014; Melander et al., 1986; Qualls-Creekmore et al., 2017; Vila-Porcile et al., 2009). We observed a decrease in galanin protein expression within GABAergic neurites, but not somata, in 4E-BP2 cKO mice, consistent with compartment-specific translation. It has been shown that both galanin KO and overexpression impair memory (Beltran-Casanueva et al., 2024; Elliott-Hunt et al., 2004; Hooversmith et al., 2019; Kinney et al., 2002; Ögren et al., 1992; Ögren et al., 2010; Stiedl et al., 2021). Our findings suggest that precise regulation of galanin translation may be important for memory formation. Future research should explore whether restoring galanin signaling can rescue the memory and behavioral deficits observed in 4E-BP2-Gad2 cKO mice.

While our findings reveal a previously underappreciated role for 4E-BP2 in inhibitory neuron function, we note several technical limitations. Both FUNCAT and vTRAP provide static snapshots of translation and may not fully reflect the dynamic temporal changes occurring during learning and memory retrieval. Also, the induction of tagged ribosomal protein L10a via AAVs may result in uneven expression across cells, as L10a exhibits biases in RNA capture (Shi et al., 2017). It remains unclear whether the isolated mRNAs represent actively translating ribosomes or stalled complexes (Zhang et al., 2020). Although *Ccl21* and *Ccr5* mRNAs showed increased in ribosomal association, corresponding protein changes were not verified in our study. Future experiments using ELISA or IF will be required to determine whether eIF4E-dependent translation activation indeed elevated chemokines and its receptors in distinct neuronal cell types.

In conclusion, 4E-BP2 in GABAergic neurons is critical for hippocampus-dependent LTM and regulates translation of specific mRNAs without globally altering global translation. Our study expands the understanding of interneuron-specific eIF4E-dependent translational control in memory and provide insights relevant to brain disorders involving inhibitory neuron dysfunction (Gkogkas et al., 2013; Hazra et al., 2013; Ribeiro et al., 2024; Sharma et al., 2021; Wiebe et al., 2019).

## Materials and Methods

### Mouse models

Mice were purchased from the Jackson Laboratory (JAX) on C57BL/6J background. Mice expressing Gad2-Cre (#010802) were backcrossed to C57BL/6J (#000446) mice for at least 10 generations. *Eif4ebp2^flx/flx^*:*Gad2-Cre* mice were generated from crossing *Eif4ebp2^flx/flx^* to *Gad2-Cre* mice as previously described and validated (Aguilar-Valles et al., 2021; Huang et al., 2024; Wiebe et al., 2019). 4E-BP2 wild-type (WT) and conditional knockout (cKO) alleles were genotyped by PCR with primers BP2-F (5′- GTCGGTCTTCTGTAGATTGTGAGT) and BP2-R (5′- GGCGATCCCTAGAAAATAAAGCCT-3′) using ear-notched samples. Cre recombinase expression was confirmed by PCR with primers Cre-F (5′- GATTGCTTATAACACCCTGTTACG-3′) and Cre-R (5′- GTAAATCAATCGATGAGTTGCTTCA-3′). Primers were purchased from Integrated DNA Technologies.

### Animal Experiments

Mice were housed in the Goodman Cancer Institute (GCI) animal facility at McGill University. All experiments involving animals were approved by the McGill University Facility Animal Care Committee (FACC) regulations, which are compliant with the guidelines established by the Canadian Council on Animal Care (CCAC). The public health service (PHS) assurance number for McGill University is F-16-00005 (A5006-01). Mice were kept in the animal facility room at a maintained temperature (20–22 °C) with a 12-hr light/dark cycle (7:00–19:00 light phase). Mice were group-housed according to sex (2-5 mice per cage). Mice had free access to food and water *ad libitum*. All experiments were conducted on male mice (3-4 months old), behavioral experiments were done during the light phase. All control experiments were performed on mice expressing Cre recombinase in GAD2^+^ neurons to normalize for any confounding effects due to the expression of Cre.

### Western blotting

Hippocampal tissues were collected and homogenized in ice-cold radioimmunoprecipitation assay (RIPA) buffer (Sigma-Aldrich, R0278) containing phosphatase inhibitor cocktail 2 (Sigma-Aldrich, P5726), phosphatase inhibitor cocktail 3 (Sigma-Aldrich, P0044), and protease inhibitor tablet (Roche, 11836170001). Lysates were incubated on ice for 30 mins before centrifuging at 16,000 × g for 20 mins at 4 °C. The supernatant was collected. Total protein concentrations were quantified using Bradford assay. Equal amounts of protein (25 μg per sample) were mixed 1:1 with 2× Laemmli loading buffer containing β-mercaptoethanol Samples were separated on 12% SDS-PAGE gels using 70V for the first hour and 100V after that. Proteins were transferred onto 0.2 μm nitrocellulose membranes (Bio-Rad, 1620112) overnight at 25 V in the cold room (4 °C). Membranes were blocked with 5% bovine serum albumin (BSA) in Tris-buffered saline (TBS, 20 mM Tris–HCl, pH 7.5, 150 mM NaCl) containing 0.1% Tween-20 (TBST) for 1 hr at room temperature (RT). Membranes were probed with diluted primary antibodies overnight at 4 °C. The following primary antibodies against the indicated proteins were used: 4E-BP2 (Cell Signaling Technology, #2845; 1:500), Neuroligin 2 (Synaptic Systems, #129203; 1:1000); anti-VGAT (Synaptic Systems, #131002; 1:1000); anti-GABABR2 (Synaptic Systems, #322203; 1:1000), anti-Gephyrin (Synaptic Systems, #147008; 1:1000); anti-Glutamate Receptor 1 (Abcam, 183797; 1:1000) and anti-GAPDH (Abcam, #9842; 1:20,000). After three washes in TBST, membranes were incubated with horseradish peroxidase (HRP)-conjugated anti-mouse or anti-rabbit secondary antibodies (Jackson ImmunoResearch, #111-035-003; 1:10,000) for 1hr at room temperature. Membranes were washed 3X with TBST and incubated for 1 min using enhanced chemiluminescence (ECL) reagents (Revvity, ORT2755 and ORT2655). Signal was visualized using Blu-Lite Ultra-High Contrast Western Blotting film (MTC Bio, A8815) and quantified using Image J software (NIH).

### Stereotactic surgeries

Mice were anesthetized under 3% isoflurane for induction of anesthesia and 2% isoflurane for maintenance through a vaporizer system. Mice were placed on a heated (37°C) surgical pad and secured in a stereotaxic frame using ear bars and a nose cone. Mice were administered carprofen (5 mg/kg, i.p.) and 0.9% saline pre-operatively to ensure analgesia and maintain hydration during the surgery process. Absence of pain reflexes was verified *via* the pedal withdrawal reflex test (paw pinch) and the corneal reflex test (eyeball response) prior to the procedure. Eye ointment (Systane, DIN02444062) was applied to each eyeball to prevent corneal drying. The scalp was shaved and disinfected with Isobetadine. A midline scalp incision (∼3 mm) was made, and the skull surface was exposed and cleaned with hydrogen peroxide (H_2_O_2_). Bregma was used as the zero-reference point for all stereotaxic coordinates. AAV injections targeting the CA1 region of the hippocampus were performed at AP: –1.9 mm, ML: ±1.0 mm, DV: –1.4 mm. A total of 1 μL of viral suspension was infused using a 33G needle. The incision was closed with Vetbond tissue adhesive (3M,1469SB), and mice were monitored post-operatively for recovery. Post-surgical monitoring was conducted daily for 3 days, with attention to general behavior and wound condition. Veterinary staff continued to monitor animals throughout the experimental timeline to ensure animal welfare.

### Behavioral testing

All behavioral assays were performed between 7:00 a.m.—3:00 p.m. under dim light. Prior to testing, mice were handled for 3 consecutive days, during which they were allowed to explore the experimenter’s hands for 1-2 mins daily. Before each test, mice were habituated to the testing room for 30 mins. Testing orders were randomized and counterbalanced across experimental groups. The investigator was blinded to genotypes during behavioral scoring.

### Morris water maze (MWM)

The MWM task was performed as previously described (Mahmood et al., 2024; Morris, 1981). The tank (diameter: 100 cm) was filled with opaque water (23°C) using non-toxic white tempera paint. A 10 cm² escape platform was submerged 2 cm below the surface, placed midway between the center and perimeter. Visual cues were fixed on the surrounding wall. Training occurred over 5 days. Mice underwent three 60-second trials per day with 30-minute intervals (Costa-Mattioli et al., 2005). Start positions varied semi-randomly among four fixed locations. Mice were allowed to search for the platform until escape or timeout. If mice failed to locate the platform within the allotted time, they were gently guided to it and allowed to remain on the platform for 10 seconds prior to removal. On day 6, the platform was removed for the probe trial. Mice were released from the quadrant opposite the former platform location and allowed to swim freely for 60 seconds. Time spent in each quadrant was recorded *via* video tracking software (HVS Image, Buckingham, UK). Visual and motor functions were controlled using a visible platform trial (1 day, 3 trials: 60 s/trial) with a flagged platform.

### Contextual fear conditioning (CFC)

The CFC test was performed as previously described (Fanselow, 2000; Mahmood et al., 2024). Mice were placed in the conditioning chamber for 2 mins of context exploration, followed by two-foot shocks (0.7 mA, 2 s each) for training. Mice remained in the chamber for 1-minute post-shock before returning to their home cages. Chambers were disinfected between sessions using odorless cleaning agents. Memory was tested 24 hrs later by reintroducing mice to the same context for 5 mins without any shocks. Freezing behavior, indicative of contextual fear memory, was scored every 5 seconds. Results are expressed as the percentage of freezing (%) epochs over total epochs.

### FUNCAT (Fluorescent Noncanonical Amino Acid Tagging)

Analysis of protein synthesis followed established protocols (Hooshmandi et al., 2024). Male mice were placed on a methionine-deficient diet (Envigo RMS Inc., TD.110208) for one week before the experiment. Body weight was monitored daily to ensure that weight loss did not exceed 20% of the initial baseline. After seven days on the diet, mice received an intraperitoneal injection of azidohomoalanine (AHA) 100 µg/g of body weight, a methionine analog used for labeling nascent proteins (Click-iT™ AHA, L-Azidohomoalanine, C10102, Thermo Fisher Scientific). Three hrs post i.p., animals were anesthetized and transcardially perfused with ice-cold phosphate-buffered saline (PBS), followed by 4% paraformaldehyde (PFA). Brain tissue was extracted, post-fixed in 4% PFA overnight at 4 °C. Coronal sections (30 μm) were obtained using a vibratome and stored in PBS. Sections were washed and incubated overnight at 4 °C in a blocking solution containing 10% normal goat serum (NGS), 0.5% Triton X-100, and 5% sucrose in PBS. The following day, Click Chemistry was employed to label AHA-incorporated proteins. Sections were incubated in a reaction cocktail consisting of 200 µM Triazole ligand, 400 µM Tris (2-carboxyethyl) phosphine (TCEP), 2 µM Alexa Fluor™ 555 alkyne (Cat. No. A20013, Thermo Fisher Scientific), and 200 µM CuSO₄ in PBS. The reaction proceeded overnight at 4 °C with gentle agitation. Following fluorescent tagging, immunofluorescence staining was carried out on the labeled sections.

### Immunofluorescence

Mice were anesthetized with 3% isoflurane delivered in oxygen. Mice were perfused with ice-cold 1x PBS then 4% PFA. Brains were collected and fixed in PFA for at least 24 hr at 4 °C. Prior to the immunofluorescence staining procedure, brain samples were transferred to a 30% sucrose solution in 1x PBS for cryoprotection. Mouse brain sections (20 µm) were collected using a Microm (HM 525/550) Cryostat. Sections were placed in 10 mM boiling sodium citrate buffer for 20 min and washed with 1x PBS three times. Sections were incubated in blocking buffer, consisting of 10% NGS and 0.5% Triton X-100 in PBS, for 1.5 hr at RT. Blocked sections were incubated in primary antibody (diluted in blocking buffer) at 4 °C overnight. Antibodies used in this study are as follows: GAD67, Millipore MAB5406, 1:5,000; 4E-BP2, Cell Signaling Technology, #2845; 1:50; Galanin, Millipore Sigma AB2233, 1:200. Sections were washed with 1x PBS three times for 5 min, then incubated in Alexa Fluor 488 goat anti-rabbit IgG (Thermo Fisher Scientific, #11034, 1:400), Alexa Fluor 594 goat anti-mouse IgG (Thermo Fisher Scientific, #10032, 1:400) and Hoechst 33342, trihydrochloride, trihydrate (Life Technologies, #3570, 1:5000) (diluted in 2% NGS) for 1 hr at RT. Brain slices were mounted with SlowFade Diamond Antifade Mountant (Thermo Fisher Scientific, S36972) and imaged with a ZEISS LSM880 laser scanning confocal microscope.

### Viral Translating Ribosome Affinity Purification (vTRAP)

TRAP-seq was performed following established protocols to optimize RNA yield and quality (Lister et al., 2024; Tavares-Ferreira et al., 2022). Hippocampus tissue was collected from male mice aged 3 months and placed in ice-cold dissection buffer containing Gibco 1× Hanks’ balanced salt solution (HBSS) (Invitrogen, 14065056), 2.5 mM 4-(2-Hydroxyethyl) piperazine-1-ethanesulfonic acid (HEPES) (Thermo Fisher Scientific, BP299100) containing sodium hydroxide (NaOH) (pH 7.4), 35 mM glucose, 5 mM MgCl₂, 100 µg/mL cycloheximide (CHX) (Sigma, C7698), and 0.2 mg/ml emetine (Med Chem, HY-B1479B). HPC samples were transferred into a bead homogenizer tube filled with ice-cold homogenization buffer, containing 20 mM HEPES-NaOH (pH 7.4), 12 mM MgCl₂, 150 mM KCl, 0.5 mM DL-Dithiothreitol (DTT) (Thermo Fisher Scientific, P2325), 1 µL/mL protease inhibitor cocktail (Roche, 11836170001), 100 µg/mL cycloheximide, 20 µg/mL emetine, 40U/mL RNasin (Promega, N2515), and 2 µL/mL TURBO DNase (Invitrogen, Invitrogen). Homogenization was carried out using a Minilys Personal Homogenizer (Bertin Technologies) with 8 × 10-second pulses at medium speed, followed by incubation on ice for 10 seconds. Post-nuclear supernatant was obtained by centrifugation at 2,000 × g for 5 mins at 4°C. The supernatant was incubated with 10% NP-40 (AG Scientific, P-1505) and 300 mM 1,2-diheptanoyl-sn-glycero-3-phosphocholine (DHPC) (Avanti Polar lipids, 850306P) for 5 mins. Post-mitochondrial supernatants were further cleared by a centrifugation step at 15,000 × g for 10 mins. An aliquot (200 µL) of the lysate was saved by adding 1 μl of TURBO DNAse (Invitrogen, AM2238) for input (IN). To isolate ribosome-bound mRNAs (immunoprecipitation, IP), cleared lysate was incubated with anti-GFP antibodies (50 µg HtzGFP-19F7 and 50 µg HtzGFP-1G8, Sloan Kettering Core Facility) pre-coupled to Protein Dynal Protein G magnetic beads (Invitrogen, 10003D) in RNAse free microcentrifuge tubes. IP was performed overnight at 4°C using an end-over-end rotator. The following day, beads were washed 4 times in a 0.35 M KCl high-salt buffer (20 mM HEPES-NaOH, pH 7.4, 12 mM MgCl₂, 300 mM KCl, 1% NP-40, and 0.5 mM DTT, 100 μg/ml CHX) to remove non-specific interactions. A total of 200 μl unbound fraction was collected for RNA extraction. Immunoprecipitated ribosome-mRNA complexes were extracted from the beads (IP), UNBOUND fraction and IN using TRIzol reagent (ration 1:3; Invitrogen, 15596026), followed by a 10-minute incubation at RT. Following incubation, samples were vortexed for 30 s and briefly centrifuged at 4 °C. Samples were placed on a magnetic rack for 1 minute to separate the beads. The RNA containing phase was carefully collected and RNA was extracted using the Direct-Zol RNA Kit (Zymo Research, R2060) according to the manufacturer’s instructions. An equal volume of 95% ethanol was added to each sample (IP, UNBOUND and IN). The mixture was transferred into Zymo-Spin columns in a collection tube. Columns were centrifuged at 16,000 x g for 30 sec and the flow through was discarded. The columns were transferred into new collection tubes. RNA was subsequently purified using the Direct-zol RNA kit, following the manufacturer’s instructions. Washing and elution steps were performed on the column matrix. RNA concentration and purity were assessed using a NanoDrop ND-1000 spectrophotometer (Thermo Fisher), and RNA integrity was verified using the 2100 Bioanalyzer (Agilent Technologies).

### Library Generation and Sequencing

Sequencing was performed on both IP ribosome-associated RNA and their corresponding IN RNA fractions. For each genotype, control and 4E-BP2 cKO, 4 biological replicates were generated, with each replicate derived from pooled 6 hippocampi of 3 mice. Total RNA was quantified, and its integrity was assessed on LabChip GXII (PerkinElmer). Ribosomal RNA (rRNA) was depleted from 70 ng of total RNA using the QIAseq FastSelect rRNA removal kit (Human/Mouse/Rat, Qiagen). cDNA synthesis was performed with the NEBNext RNA First Strand Synthesis and NEBNext Ultra Directional RNA Second Strand Synthesis Modules. Library construction continued using the NEBNext Ultra II DNA Library Prep Kit for Illumina (New England Biolabs). Adapter oligonucleotides and PCR primers were obtained from New England Biolabs. Libraries were quantified using the KAPA Library Quantification Kit (KAPA Biosystems) and average library fragment size was determined using a LabChip GX system (PerkinElmer). Libraries were normalized, pooled, and the DNA was denatured with 0.05M NaOH, followed by neutralization with HT1 hybridization buffer. Sequencing was conducted on an Illumina NovaSeq S4 platform using Xp protocol and 2 × 100 cycle paired-end mode, in accordance with the manufacturer’s instructions. PhiX control DNA was added at 1% to monitor sequencing quality. Base calling was performed using Real-Time Analysis (RTA) software v3.4.4, and demultiplexing was conducted with bcl2fastq2 v2.20 to generate FASTQ files (Tavares-Ferreira et al., 2022).

### Bioinformatics and Statistical Analysis

#### Read Alignment and Transcript Quantification

Initial quality control of raw sequencing data was conducted using FastQC (Babraham Bioinformatics, https://www.bioinformatics.babraham.ac.uk/projects/fastqc/), assessing per-base quality, sequence content, and duplication levels. Adapter trimming and removal of low-quality bases (12 bases per read) were performed. Processed reads were aligned to the *Mus musculus* reference genome (GRCm39, Ensembl release M31, primary assembly) using STAR (v2.7.6) (Dobin et al., 2012). Following alignment, deduplication was carried out using Sambamba (v0.8.2) (Tarasov et al., 2015). Transcript abundance was subsequently quantified with StringTie (v2.2.1), yielding transcript-per-million (TPM) values for each gene across all samples (Pertea et al., 2015).

#### Expression Data Normalization and Order statistics

Subsequent analysis of TRAP datasets followed established protocols (Lister et al., 2024; Sanz et al., 2009; Tavares-Ferreira et al., 2022). For each RNA-seq and TRAP-seq sample, expression consistency was assessed by calculating percentile ranks of TPM values for all coding genes. Genes consistently expressed above the 30^th^ percentile across all IN samples (14,804 genes) were considered robustly expressed and retained for further analysis. Genes consistently detected above the 15^th^ percentile in IP samples (12,917 genes) were selected for downstream analysis. These numbers are consistent with previous TRAP studies from mouse samples (Tavares-Ferreira et al., 2022). To correct for sequencing depth and identify actively translated mRNAs, percentile ranks of TPM values for consistently expressed genes in the IP fractions were recalculated and quantile normalized prior to comparative analysis.

#### Differential Expression Analysis

Differential expression (DE) analysis was performed using previously established approaches (Sanz et al., 2009; Tavares-Ferreira et al., 2022). For each mRNA, the log_2_ fold change (Log_2_FC) between control and 4E-BP2 conditional KO was computed using normalized TPM values. mRNAs were evaluated in both the transcriptome (IN) and translatome (IP) datasets. To assess reproducibility and effect size, the Strictly Standardized Mean Difference (SSMD) was calculated for each gene across biological replicates. SSMD quantifies the signal-to-noise ratio and was complemented by the Bhattacharyya distance which measures the degree of overlap between expression distributions. In our analysis, modified Bhattacharyya coefficient that ranges between 0 (for completely identical distributions) and +1 or −1 (for totally nonoverlapping distributions, sign defined by the log-fold change value) was used. Genes were considered differentially expressed if the absolute SSMD exceeded 0.9, the absolute BC surpassed 0.5, and the absolute log_2_FC was >1.33 (|Log_2_FC |>0.41) (Tavares-Ferreira et al., 2022). All computational analyses and data visualization were performed using Python (v3.7) and Anaconda.

#### Gene ontology analysis

GO enrichment analysis of differentially expressed mRNAs in IP was performed using Enrichr (http://amp.pharm.mssm.edu/Enrichr/) (Kuleshov et al., 2016). Pathways with p< 0.05 were considered significantly enriched by differentially expressed mRNAs (minimum of 3 associated genes).

#### Statistical analysis

Data analysis was performed using GraphPad Prism 10. Bar plots display means with SEM. One-sample t-tests were used to compare group means to hypothetical values. Depending on the experimental design, data were analyzed using two-tailed unpaired Student’s t-tests, Welch’s correction (for unequal variances), two-way ANOVA, and post hoc comparisons (Bonferroni’s test) as appropriate. Repeated measures were assessed *via* two-way repeated measures ANOVA or mixed-effects models. Results are presented as mean ± standard error of the mean (SEM). A p-value < 0.05 was considered statistically significant (n.s.: p > 0.05; *p ≤ 0.05; **p ≤ 0.01; ***p ≤ 0.001; ****p ≤ 0.0001).

## Abbreviations

4E-BP: eukaryotic initiation factor 4E-binding protein
AAV: adeno-associated virus
AD: Alzheimer’s disease
cKO: conditional knockout
eIF4E: eukaryotic initiation factor 4E
eIF4G: eukaryotic initiation factor 4G
ERK: extracellular signal-regulated kinase
FMRP: fragile X messenger ribonucleoprotein
GABA: gamma-aminobutyric acid
GAD2: glutamate decarboxylase 2
LTM: long-term memory
MNK: mitogen-activated protein kinase-interacting kinase
mTORC1: mechanistic/mammalian target of rapamycin complex 1
mTORC2: mechanistic/mammalian target of rapamycin complex 2
PC: Purkinje cell
PI3K: phosphoinositide 3-kinase
RPTOR: regulatory associated protein of mTOR
S6K: ribosomal protein S6 kinase
SST: somatostatin
TRAP: translating ribosome affinity purification
TSC: tuberous sclerosis complex

## Data availability

The research data are available from the corresponding author on reasonable request.

## Ethics approval

All animal procedures and experiments were performed in accordance with the McGill University animal care committee regulations.

## Acknowledgements

We thank Karim Nader (McGill University) for providing access to behavioral testing facilities and Wayne Sossin (McGill University) for critical comments. This study was supported by Canadian Institutes of Health Research (CIHR) foundation grant (FDN-148423) and Natural Sciences and Engineering Research Council of Canada (NSERC) Discovery Grants (RGPIN-2021-02764). N.M. is supported by a postdoctoral fellowship from the Goodman-Kahvejian family.

## Author contribution

Z.H. designed and performed experiments, interpreted the data, and wrote the manuscript. N.M. and K.P. performed behavioral experiments and analyzed data. K.L. and M.H. helped with FUNCAT and vTRAP experiments. N.I., D.T.F. and M.A. analyzed the vTRAP data. S.W. and A.K. provided conceptual support. N.S. provided supervision and secured funding. All authors contributed to editing the manuscript and approved the final version.

## Consent for publication

All authors have given their consent for publication.

