## Supplemental figures for "4E-BP2-dependent translational control in GABAergic interneurons is required for long-term memory"

**is required for long-term memory**

a. Department of Biochemistry, McGill University, McIntyre Medical Building, 3655 Promenade Sir William Osler, Montréal, QC, Canada H3G 1Y6.
b. Goodman Cancer Institute, 1160 Pine Avenue West, Montréal, QC, Canada H3A 1A3.
c. Department of Anesthesia and Faculty of Dental Medicine and Oral Health Sciences, McGill University, Montréal, QC, Canada H4A 3J1.
d. School of Behavioral and Brain Sciences and Center for Advanced Pain Studies, University of Texas at Dallas, Dallas, United States, 75080.
e. Current address: Cancer Research Malaysia, Level 1, Subang Jaya Medical Centre South Tower, No. 1, Jalan SS 12/1A, 47500 Subang Jaya, Selangor, Malaysia.
f. Integrated Program in Neuroscience, McGill University, Montréal Neurological Institute, 3801 University Street, Montréal, QC, Canada H3A 2B4.

**Figure legends**

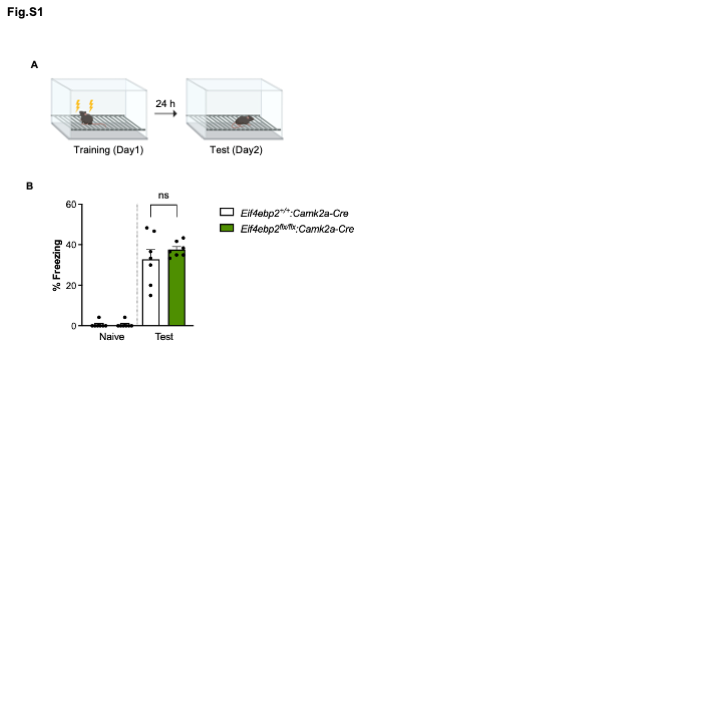

**Figure S1. Conditional deletion of 4E-BP2 in CAMK2A+ excitatory neurons does not impact contextual fear memory.**

(A) Diagram of the weak contextual fear conditioning (CFC) paradigm (2-foot shocks, 0.7 mA, 2 s duration). (B) Long-term memory (LTM) assessed as freezing behavior during a 5-minute test, 24 hours post-training. Behavioral experiments were conducted in the same cohort of mice: control (n = 7) vs. 4E-BP2-CaMK2A cKO (n = 7). Data represent individual mice and are expressed as mean ± SEM. *P < 0.05, n.s., not significant.

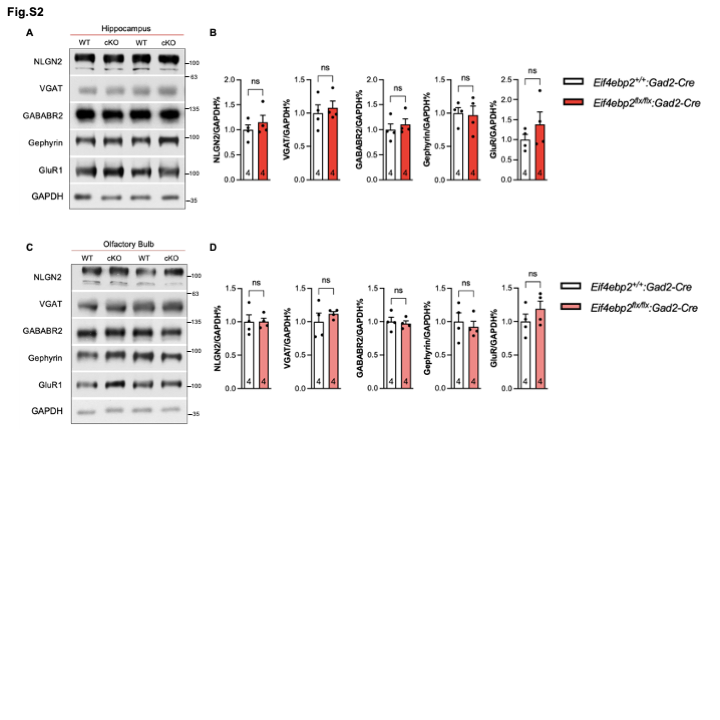

**Figure S2. 4E-BP2 deletion in GABAergic neurons does not affect the expression level of key synaptic proteins in the hippocampus and Olfactory bulbs.**

(A) Representative western blot images showing protein expression in hippocampal tissue from adult WT and 4E-BP2-GAD2 cKO mice. Western blot analysis was performed to detect NLGN2, VGAT, GABABR2, gephyrin, and GluR1. (B) Quantification of band intensities for NLGN2, VGAT, GABABR2, gephyrin, and GluR1. Densitometry was performed using ImageJ, with values normalized to GAPDH. (C) Representative western blot images showing protein expression in olfactory bulb tissue from adult WT and 4E-BP2-GAD2 cKO mice. Western blot analysis was performed to detect NLGN2, VGAT, GABABR2, gephyrin, GluR1. GAPDH served as a loading control. Data represent individual mice and are expressed as mean ± SEM. *P < 0.05, n.s., not significant. (D) Quantification of band intensities for NLGN2, VGAT, GABABR2, gephyrin, and GluR1. Densitometry was performed using ImageJ, with values normalized to GAPDH. Statistical analysis was conducted using student t-test. Data represent individual mice and are expressed as mean ± SEM. *P < 0.05, n.s., not significant.

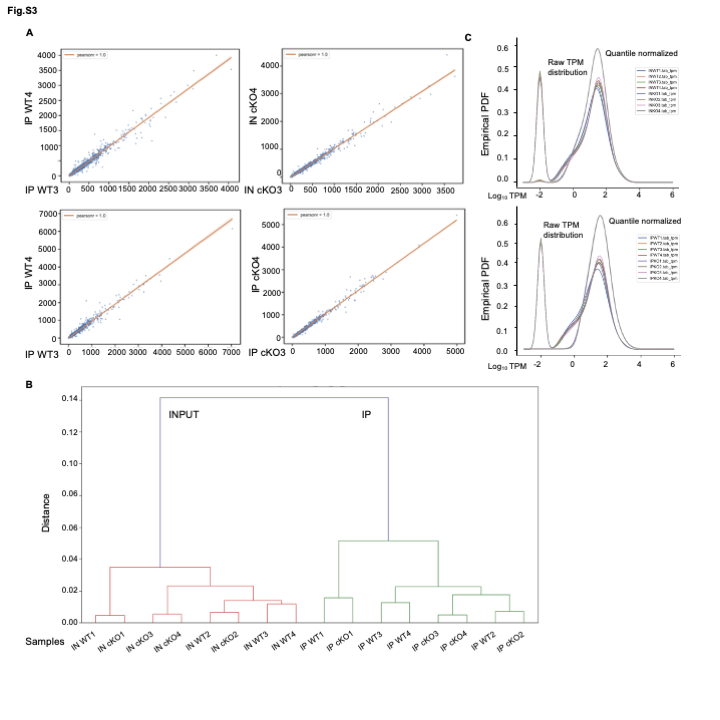

**Figure S3**. **vTRAP-Seq: quality control and data analysis**

(A) Representative correlation plots demonstrate strong linear relationships in gene expression (TPM) across biological replicates for both input (IN) and immunoprecipitated (IP) fractions. Data are shown for four replicates per genotype and condition, indicating high reproducibility within each group. (B) Hierarchical clustering analysis of the correlation coefficient show clear separation between TRAP-Seq (IP) and bulk RNA-Seq (IN). (C) Empirically estimated probability density plots illustrate the distributions of raw TPM values and quantile normalized TPM (qTPM) values for both IN and IP fractions across all samples. PDF, probability density function; TPM, transcripts per million; TRAP-Seq, translating ribosome affinity purification sequencing.

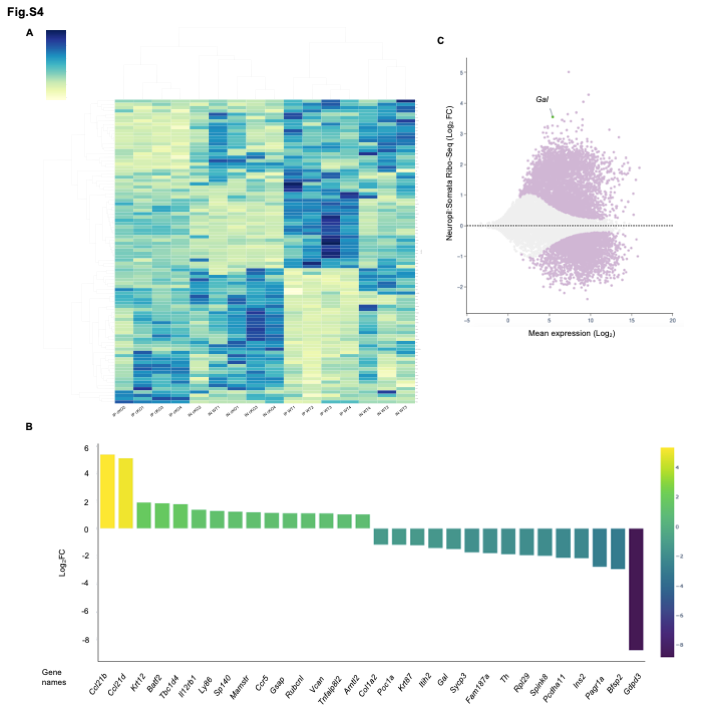

**Figure S4**. **vTRAP-Seq differentially translated mRNAs in WT and cKO hippocampus.**

(A) Heatmap shows the z scores of the Differentially expressed mRNAs in TRAP-Seq (IP) samples. Labels represent biological replicate number. (B) Top 15 most significantly upregulated and downregulated genes in GABAergic neurons from 4E-BP2-Gad2 cKO mice. Log_2_FC, Log_2_ fold change. (C) Comparison of Ribo-Seq data from neuropil versus somatic compartments for the gene Galanin. Data derived from Glock et al. (2021), The mRNA translation landscape in the synaptic neuropil, Max Planck Institute for Brain Research. *[Accessed May 2025 from* [*https://public.brain.mpg.de/dashapps/localseq/*](https://public.brain.mpg.de/dashapps/localseq/)*].*
